# Michigan ZoomIN: validating crowd-sourcing to identify mammals from camera surveys

**DOI:** 10.1101/2020.06.09.143180

**Authors:** Gabriel I. Gadsden, Rumaan Malhotra, Justin Schell, Tiffany Carey, Nyeema C. Harris

## Abstract

Camera trap studies have become a popular medium to assess many ecological phenomena including population dynamics, patterns of biodiversity, and monitoring of endangered species. In conjunction with the benefit to scientists, camera traps present an unprecedented opportunity to involve the public in scientific research via image classifications. However, this engagement strategy comes with a myriad of complications. Volunteers vary in their familiarity with wildlife, and thus, the accuracy of user-derived classifications may be biased by the commonness or popularity of species and user-experience. From an extensive multi-site camera trap study across Michigan U.S.A, images were compiled and identified through a public science platform called *Michigan ZoomIN*. We aggregated responses from 15 independent users per image using multiple consensus methods to assess accuracy by comparing to species identification completed by wildlife experts. We also evaluated how different factors including consensus algorithms, study area, wildlife species, user support, and camera type influenced the accuracy of user-derived classifications. Overall accuracy of user-derived classification was 97%; although, several canid (e.g., *Canis lupus, Vulpes vulpes*) and mustelid (e.g., *Neovison vison*) species were repeatedly difficult to identify by users and had lower accuracy. When validating user-derived classification, we found that study area, consensus method, and user support best explained accuracy. To continue to overcome stigma associated with data from untrained participants, we demonstrated their value by showing the accuracy from volunteers was comparable to experts when classifying North American mammals. Our hierarchical workflow that integrated multiple consensus methods lead to more image classifications without extensive training and even when the expertise of the volunteer was unknown. Ultimately, adopting such an approach can harness broader participation, expedite future camera trap data synthesis, and improve allocation of resources by scholars to enhance performance of public participants and increase accuracy of user-derived data.

---

Science is a public good. In recent years, an interest in broadening participation in science has resulted in a wave of public (participatory, community, or citizen) science projects initiated by scholars (Dickinson et al. 2010, Theobald et al. 2015). The seemingly unlikely partnership between the public and scientists creates dialogue that can aid in demystifying the scientific process, facilitate exposure to the natural world, and yield reciprocal benefits (Bonney et al. 2016, Schuttler et al. 2018b). Public participation in ecology is particularly salient due to humans being a dominant force in natural systems with increasing pressures stemming from land conversion, urban sprawl, and over-exploitation (Raudsepp-Hearne et al. 2010, Halpern et al. 2015, Young et al. 2016). The public can gather large amounts of data during a person’s everyday activity to monitor biodiversity and environmental change (Trumbull et al. 2000). For example, someone interested in how urbanization affects species can assist in collecting bird skyscraper collisions or simply take images with a smartphone and document sightings of particular organisms (Ries and Oberhauser 2015, Shipley et al. 2018).

Digital technology, including cameras, is emerging as an important tool for participatory science given the multitude of highly accessible platforms available to share data. As avid users of technology, engaged youth have the potential to amplify positive effects of participatory science through intergenerational learning (Arts et al. 2015, Peterson et al. 2019). Hence, incorporating images garnered from camera surveys are now commonplace for public science programs (e.g., Schuttler et al. 2018a). Camera surveys can answer a plethora of ecological, behavioral, and even evolutionary questions across spatial and temporal scales, in a variety of environments, and on a diversity of species (Burton et al. 2015, McShea et al. 2015, Steenweg et al. 2017).

Although crowdsourcing has gained popularity, image identification from wildlife surveys by communities of non-scientists is fraught with challenges (Hochachka et al. 2012). Researchers not only have to develop the infrastructure to facilitate easy access, but also construct a useful workflow to aggregate and structure responses into manageable data (Waetjen and Shilling 2017, Egerer et al. 2019, Parrish et al. 2019). Chief amongst the rising concerns is that public-generated data are susceptible to biases and inaccuracy of responses that can compromise usability and hinder broader implementation in future scientific research (Dickinson et al. 2010, Hochachka et al. 2012, Brown et al. 2018). By design, researchers are often not able to control for who contributes to citizen science projects, resulting in heterogeneity in user expertise. Projects also vary in the extensiveness of aids such as sample photos or comparison charts for similar species that could alter a user’s capacity to accurately classify images (Skarlatidou et al. 2019). Through targeted outreach, engagement, and education strategies, some projects are able to target specific audiences (e.g., K-12) and thoroughly train volunteers; though, this is not always feasible and requires significant resources. Varying methods of image classification, and the implementation of automated and crowd-sourcing aids in classifying images to discern patterns from large amounts of biological information can be helpful in reducing processing time (Maes et al. 2015, Norouzzadeh et al. 2018). Standardized guidelines for processing and evaluating user-derived data will allow for scientists to better evaluate their utility and leverage public science to tackle large data sets regardless of the research system (Freitag and Pfeffer 2013, Maes et al. 2015, Egerer et al. 2019, Locke et al. 2019).

Overall, findings have been inconsistent concerning reliability of user-derived data for scientific research, requiring additional scrutiny based on project workflows and inclusion of expert validation (Danielsen et al. 2005, Crall et al. 2011). Katrak-Adefowora et al. (2020) found that training allowed participants even with no biology background to correctly identify images with comparable accuracy to biology experts. Other studies found that there was high agreement among classifying individuals, indicating that user expertise was not significant in correct classifications, but animal size and distinct markings had a greater impact on accuracy (Swanson et al. 2016, Potter et al. 2019). Still other studies have found that environmental, species-specific behaviors, and land use affect classification accuracy (Broeckhoven and le Fras Nortier Mouton 2015, Meek et al. 2016). Generally, existing studies on validation of user derived data fail to explain factors contributing to accuracy, occur outside a North American context, or employ a single metric for deriving accuracy from aggregated responses. Another consideration that remains unexplored for camera surveys is whether data generated from public participants are comparable across geographies. Potential differences in species composition and image quality from environmental conditions such as weather may influence how the public classifies images, which may affect accuracy (Meek et al. 2015, Steger et al. 2017).

Here, we used public generated image identifications from an extensive multi-site, multi-season camera trap survey throughout the state of Michigan. Specifically, we created a public science online platform called *Michigan ZoomIN (MZI)* for image processing and developed a science communication strategy to solicit volunteers. We then evaluated user responses by comparing various consensus methods to obtain a converged species identification as well as examined factors that influenced the accuracy of users. We hypothesized that common species will be classified with higher accuracy while physically similar or rare species would have lower user accuracy and reach consensus using a relaxed consensus method. Using a hierarchical approach to assess user responses, we hypothesized high agreement would positively correlate with accuracy for all images regardless of consensus method and species characteristics. We also hypothesized that camera type would influence user accuracy with images from cameras that used a flash at night having higher accuracy. We hypothesized that the study area would affect accuracy, as sampling locations varied in vegetation and environmental conditions that could distort images and make classifying harder. We contribute three novel insights to research involving image identification from crowdsourcing by: 1) determining whether hierarchical consensus criteria can mitigate potential bias from volunteers with varying levels of expertise, 2) assessing multiple consensus thresholds to compare accuracy among species, and 3) identifying various factors that can diminish accuracy. Results from our work also have relevance in minimizing workload in classifying for researchers, as well as providing a workflow for assessing overall accuracy of user-derived classifications for other camera studies.

## STUDY AREA

Throughout 2015—2017, the Applied Wildlife Ecology (AWE) lab at the University of Michigan implemented camera trap surveys to investigate questions of community ecology within the carnivore guild of Michigan. We deployed 273 remotely-triggered Reconyx© cameras (PC 850, PC 900, PC 850C, PC 900C) across three study areas that had distinct environmental and habitat attributes from north to south: the Huron Mountain Club (HMC; Marquette County), University of Michigan Biological Station (UMBS; Pellston County), and the Shiawassee National Wildlife Refuge (SNWR; Saginaw County). The Huron Mountain Club was our most ecologically diverse site, comprised privately-owned mixed secondary and old growth forest along the southern coast of Lake Superior. The University of Michigan Biological Station, a research station, comprised a mix of deciduous and coniferous forest between Douglas and Burt Lakes in the northern tip of Michigan’s Lower Peninsula. The Shiawassee National Wildlife Refuge, a floodplain of Saginaw County, comprised marsh, bottomland hardwood forest, and grasslands abutting urban development and agriculture managed by the U.S. Fish & Wildlife Service. At each site, unbaited cameras were placed 0.5—1.0 m off the ground and affixed to trees > 0.5m diameter. Camera settings included: high sensitivity, one-second lapse between a sequence of three pictures in a trigger, and a 15-second quiet period between triggers. Our survey efforts across 32,542 trap nights resulted in 10,199 of camera trap images that required accurate species identifications.

## METHODS

### Description of *Michigan ZoomIN*

We developed a public science website called *Michigan ZoomIN (MZI)* on the Zooniverse Project Builder platform where we uploaded images and collated user responses over multiple seasons (Figure 1). From its origins in 2007 with the Galaxy Zoo project, which asked volunteers to classify shapes of galaxies, the Zooniverse crowdsourcing platform has created and hosted more than 200 different projects from disciplines as diverse as astronomy, ecology, physics, history, and more to promote people-powered research (Lintott 2019; Zooniverse.org). Through both a coordinated outreach program throughout Michigan and the existing Zooniverse community, 645,000 classifications were completed over one year by 3,950 registered users (unregistered users also contributed but were not trackable). There were 39 possible classifications ranging from the focal carnivores in our empirical field research (e.g., coyote *Canis latrans*, raccoon *Procyon lotor*, American black bear *Ursus americianus*, red fox *Vulpes vulpes*), and their prey (e.g., white-tailed deer *Odocoileus virginianus*, squirrel *Sciurus carolinensis*, opossum *Didelphis virginiana*, cottontail rabbit *Sylvilagus floridanus*, porcupine *Erethizon dorsatum*, small mammal). Users were given written descriptions of species as well as photos to aid in classifying species from survey images (Figure 1 – label 7). We also incorporated the “looks like” tool in MZI, which allows volunteers to narrow possible classifications based on color, shape, and body size (Figure 1 – label 5). We did not include ‘unknown’ as a possible choice for selection to promote more discrimination and an “educated” guess. Users were also provided a field guide with descriptions of common ecological tenets such as competition and dietary separation, and additional images and information useful for classification of species (Figure 1 – label 6). The field guide resource was included to provide more education to improve understanding and stimulate greater enthusiasm particularly for younger audiences. Images were retired from viewing once 15 unique users classified the image.

**Figure 1.**
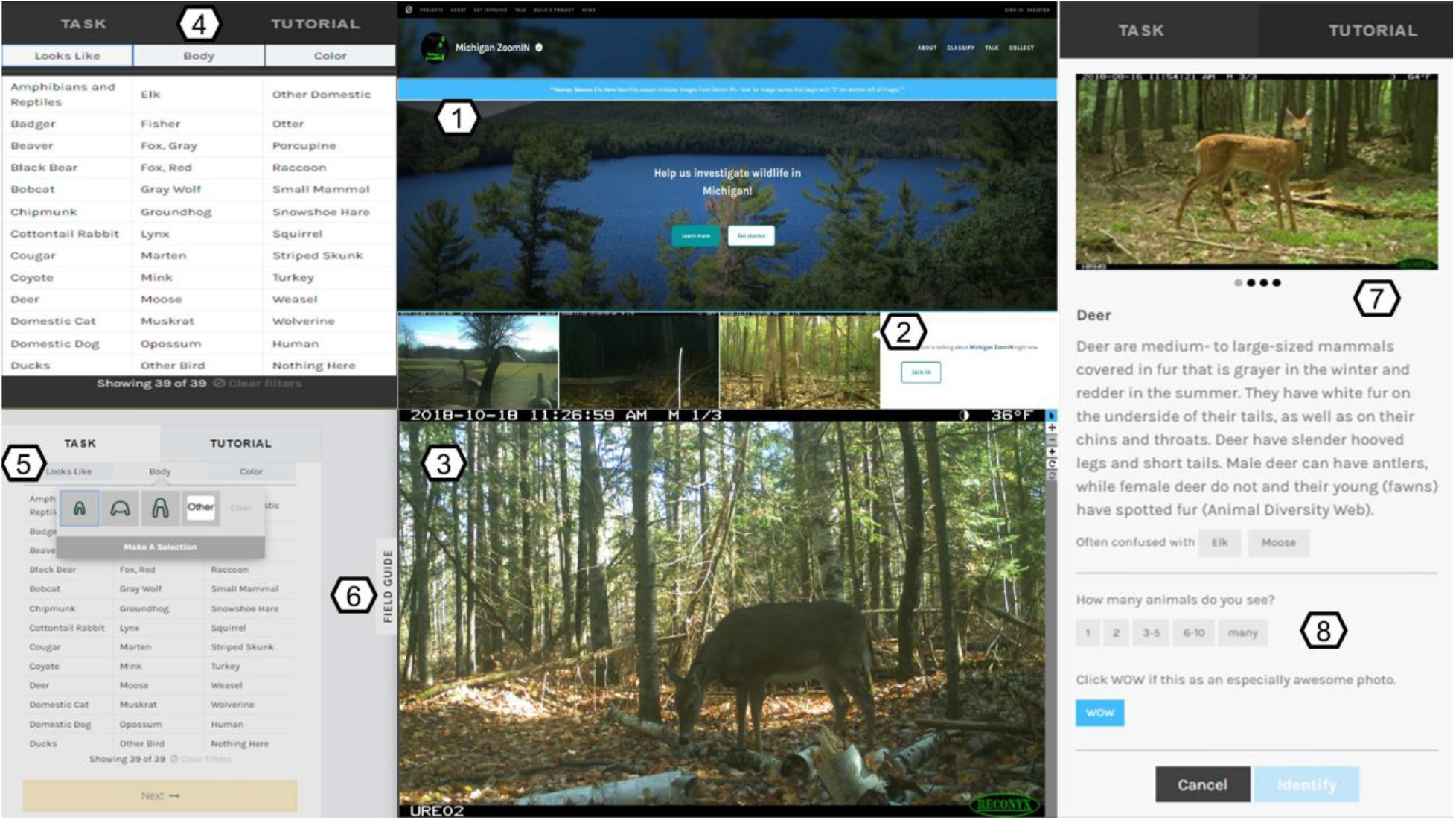
*Michigan ZoomIN* interface for public user classifications of mammals from camera surveys throughout Michigan. 1) welcome screen with tutorial, 2) sample images active on talk board from public comments, 3) image for user identification, 4) list of species to select for classification, 5) filter features to narrow species selections, 6) field guide to provide additional information, 7) reference photo of user selection to compare to image, and 8) additional questions to provide more information concerning the image.

### Data Aggregation

To produce a final identification for multiple user responses, we aggregated classifications into a single image species identification using various consensus approaches (Figure 2). Images were considered to have reached consensus if they matched the criteria of any one of four consensus methods: agreement, Kappa, Kappa and evenness, or gold standard. 1) Agreement: if all 15 users agreed on the species identification for an image. 2) Kappa: A Fleiss Kappa statistic (modified from Cohen’s Kappa, for more than 2 users) above 0.8 designating a substantial strength of agreement (Landis and Koch 1977). 3) Kappa and Evenness: A Fleiss Kappa above 0.6 (lower than the recommendations of Landis and Koch (1977)), and evenness below 0.5. Pielou’s evenness is an index that measures the proportion of votes given to a particular species in comparison to the number of species picked for the image in question. The scale runs from 0—1, with 0 indicating low evenness and higher accuracy (Pielou 1966, Swanson et al. 2015). 4) Gold Standard: At least one of the users classifying the image was from our research team and thus categorized as an expert with a gold standard tag. If the expert was unsure of classification, they removed the gold tag prior to identification submission. Images that did not reach any consensus method were classified as Majority and later verified by a project researcher (Figure 2). The following process allowed us to test a hierarchical approach that ranged from rigorous (100% user agreement) to more relaxed criteria of user agreement. We specifically tested Kappa and Kappa and Evenness as metrics that allow progressively lower levels of user agreement to account for the differing levels of experience inherent to a diverse group of users, while producing comparable accuracy to stricter consensus criteria (100% user agreement amongst all 15 users).

**Figure 2.**
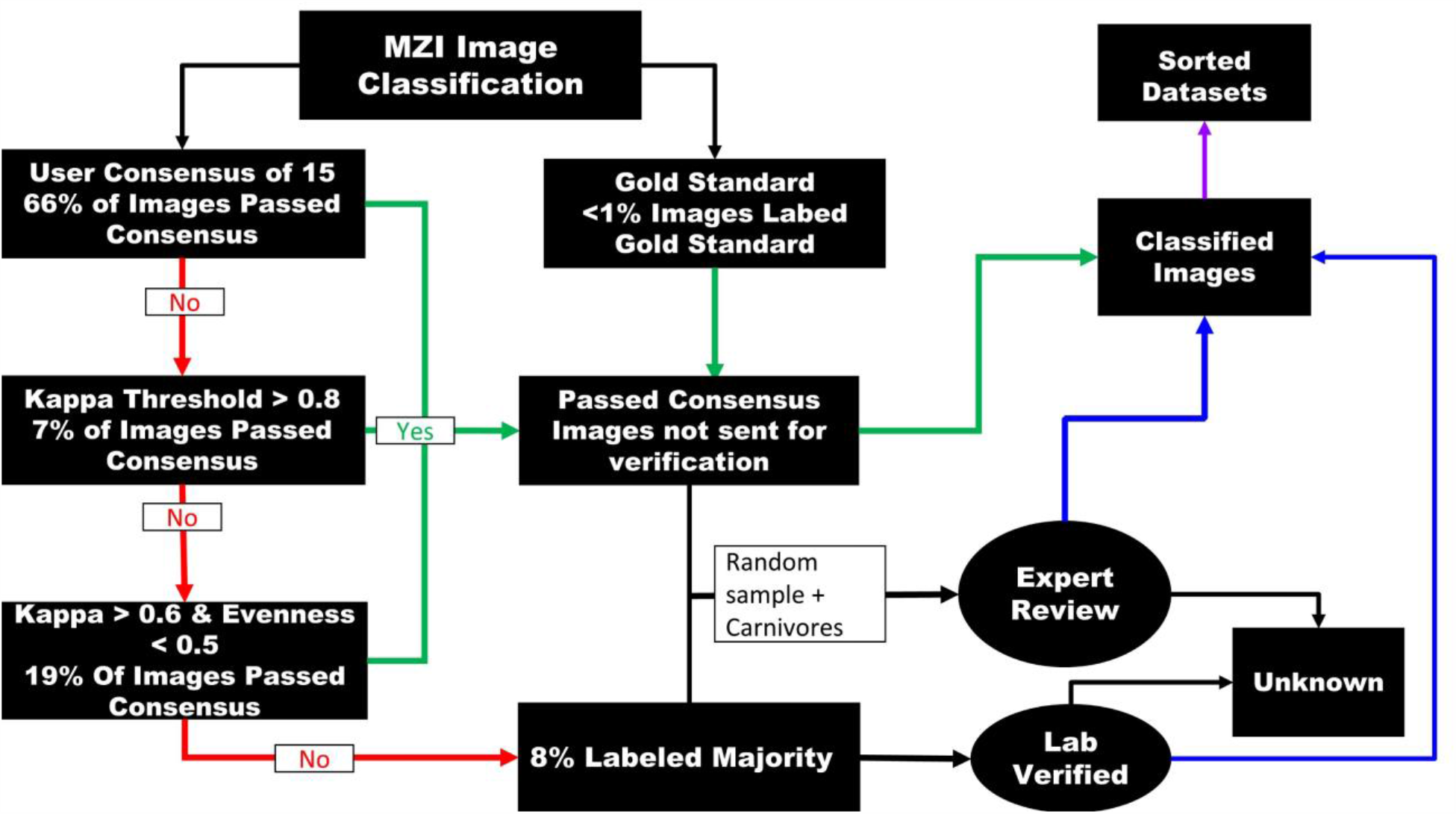
*Michigan ZoomIN* workflow from user-derived input to consensus assessment of aggregated responses to expert validation necessary to create a final dataset ready for analysis of classifying images. Images classified by an expert are tagged with gold standard which immediately passed consensus and sent to the final data set. All other images went through a hierarchical process with 15 different users classifying each image. Images that have complete agreement among users went to the final dataset. If there is not unanimous agreement, different algorithms were calculated from user responses to determine if they meet the Kappa threshold and if not the kappa evenness threshold. Images that do not meet any consensus went into an expert review file, which was then classified by a research team member to determine species.

### Expert Review

To test the efficacy of the consensus algorithm, researchers verified a random sample of 5,000 images to compare against the user-generated identifications. In addition, all images that had a carnivore identification (with the exception of raccoons) generated by the consensus algorithm were also reviewed, for a total review dataset of n = 5,734. Beyond the initial random sample of 5,000 images, additional raccoon images were not included for expert verification due to the assumption that they did not fit the rarity criterion and were well represented in the random sample.

### Evaluation of Accuracy

As a rough measure of the accuracy for each of the consensus methods used in the consensus algorithm, we calculated the proportion of correct answers for each method. For those images that did not meet any of the criteria to produce consensus identification, we calculated the proportion correctly identified among the 15 user classifications. For hypothesis testing, we first constructed a generalized linear model to evaluate factors influencing accuracy. Aggregated user responses from the consensus output were compared to the expert dataset to generate a binary output of 1(correct) or 0 (incorrect). For comparison against images that were classified by consensus, images that did not meet a consensus criterion were classified under the consensus type as majority. Majority represented the most widely chosen species by public users when consensus was not reached by other methods. We included: consensus methods, species, study area, camera model, and percent user agreement as fixed effects in the model. Species was used to better understand if animal similarity or rarity of a species affected the accuracy. We used two different camera models in our study. Infrared cameras provide black and white images in low light settings, while white flash cameras produce color images, even at night. Percent user agreement represented a measure on how well users agreed on a classification and corresponding difficulty depending on species. We conducted model selection using Akaike information criterion (AIC). All models < 3 ΔAIC were considered top models and beta coefficients for variables from top selected models were then used to test the following hypotheses:

1. Species that have a high degree of similarity will be harder to identify, which will negatively affect accuracy;
2. Species accuracy will be greater for species that are more common and have greater abundance indexed by trigger activity;
3. Stronger similarity in user agreement will correlate to overall higher accuracy;
4. White flash cameras will yield higher accuracy;
5. Accuracy will vary across study areas with the site closest to human habitation yielding higher accuracy.

All model construction and selection were conducted using the lme4 package in R (version 3.4.4).

## RESULTS

From our public science project, *Michigan ZoomIN* involved 3,950 registered users that were involved in classifying 10,199 image sequences across three study areas. (HMC: 1,630; UMBS: 2,485; SNWR: 6,084). Each sequence consisted of three images and was reviewed by 15 unique users. The most common species were raccoon and white-tailed deer with 5,594 (51%) and 2,592 (14%) classifications, respectively. Most images (95%) reached a consensus method to converge on a specific species identification. The remaining 5% remained unidentifiable even after being evaluated by a team researcher (Figure 2). Two-thirds of images were classified from complete agreement among all 15 users. Raccoon, deer, and striped skunk (*Mephitis mephitis*) were the only species to have higher than 50% of their images reach agreement (Figure 3). Kappa evenness was the second most common consensus method, comprising 19% of images. Majority, yielded completion of 10%, and majority plus gold standard comprised < 1% of images. Groundhog (*Marmota monax*), fisher (*Pekania pennanti*), mink (*Neovison vison*), muskrat (*Ondatra zibethicus*), and snowshoe hare (*Lepus americanus*) had 100% of their images go to majority, and all these species represented the fewest captures in the dataset (Figure 3).

**Figure 3.**
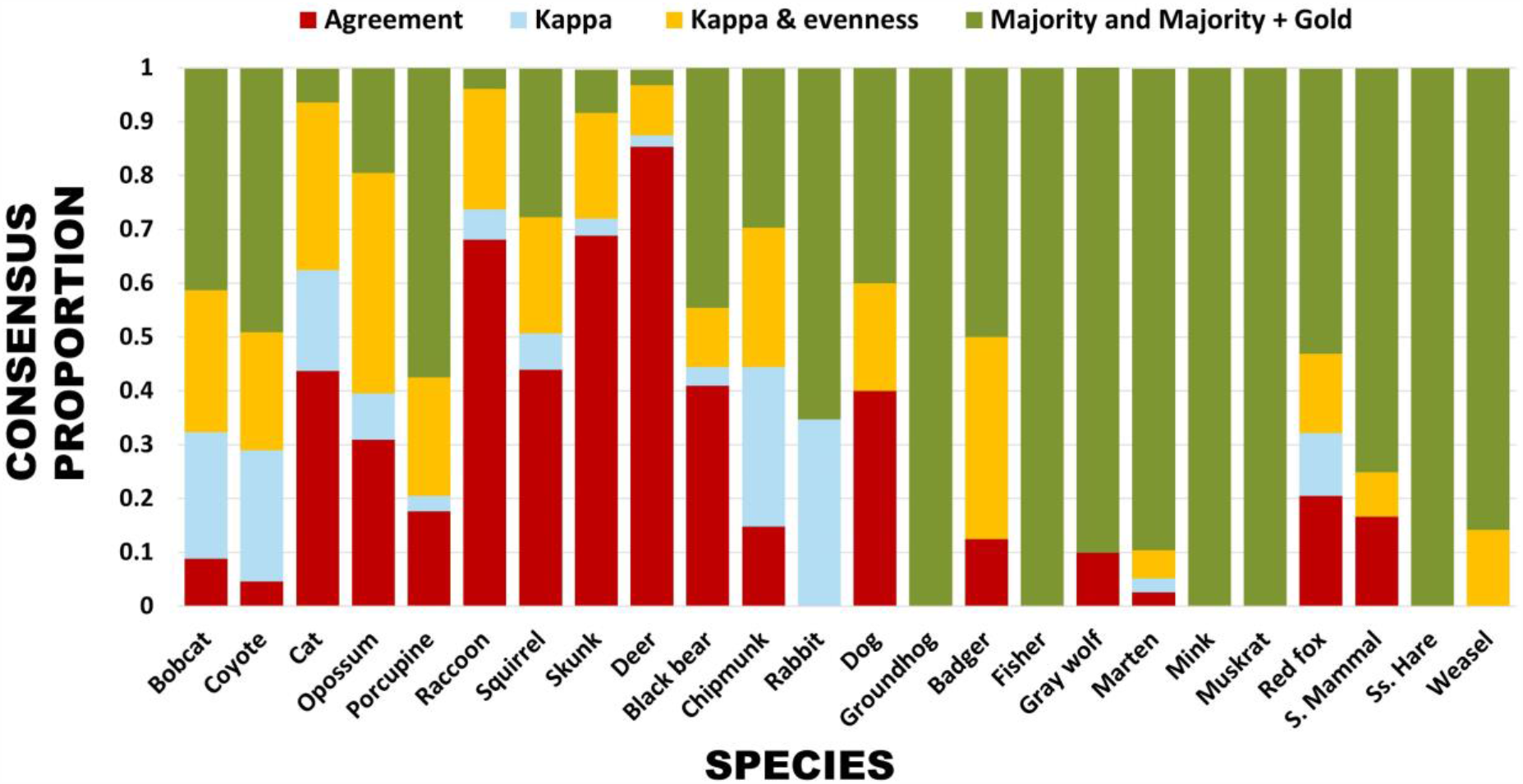
Distribution of images for each species reaching consensus across various criteria. Agreement resulted when all 15 users agreed on the species identification. Kappa was based on a Fleiss Kappa statistic above 0.8, while Kappa and evenness used Fleiss Kappa statistic above 0.6 combined with Pielou’s evenness below 0.5. Majority represented the most widely chosen species by public users when consensus not reached by other methods. Gold standard indicated when an expert contributed to the image classification, but only represented <1% of data.

We built generalized linear models to evaluate factors that influenced the accuracy of species classifications. The top model included: consensus method, percent user agreement, and study area (Table 1). Less strict consensus criteria were only marginally worse except for majority: Kappa (β = −1.61; *p* = 0.014), kappa evenness (β = −1.162; *p* = 0.058), and majority (β = −3.355; *p* <0.0001). Percent user agreement (β =5.328; *p* < 0.0001) was strongly associated with higher accuracy. Contrary to expectations, the camera model did not influence accuracy. As previously stated, consensus of 100% user agreement was the most common method in which images reached the final dataset. While most species were represented in images that reached consensus, species varied in which consensus method was most effective for identification. Indeed, the wide geographic coverage of the camera surveys presented the possibility for differences in accuracy based on study area. With a 0.1 p-value threshold, images from SNWR were significant in identification accuracy compared to other study areas (β= 5.047; p-value = 0.059). In contrast, images from HMC, the most remote site with the highest species diversity located in the Upper Peninsula of Michigan, had the lowest accuracy of species classification.

**Table 1.** Top models presented with ΔAIC < 3 from model selection analysis for accuracy assessment from comparison between user derived classification and expert validation.

| Model | df | logLik | $\Delta AIC$ | W |
| --- | --- | --- | --- | --- |
| Consensus method + Percent user agreement + Study area | 8 | -570.335 | 0.00 | 0.493 |
| Consensus method + Percent user agreement + Study area + Camera model | 9 | -570.327 | 1.99 | 0.182 |
| Consensus method + Percent user agreement + Species | 30 | -549.265 | 2.03 | 0.179 |
| Consensus method + Percent user agreement + Species + Study Area | 32 | -547.451 | 2.43 | 0.146 |

Notably, the public was 99% accurate in identifying false positives, incidental triggers from wind or vegetation movement. Small mammals were often confused as partial images of cottontail rabbits, or mistakenly missed and labeled as nothing there. Though many species had high accuracy (>90%, Figure 4), accuracy was statistically significant for squirrels (β= 2.180; *p* = 0.041) and porcupines (β= 2.299; *p* = 0.042) likely because of the ubiquity of squirrels and the uniqueness of porcupines. Classifying animals incorrectly were common only for particular species such as mustelids and coyotes. The high false positives were due to similarities to other species based upon body size, color, and activity patterns. Mustelids had the lowest overall accuracy (Figure 4); arguably, because the family comprises species with similar anatomical features (e.g., marten *Martes americana*; long-tailed weasel *Mustela frenata*; fisher; mink) that expectedly would prove difficult to discern for non-experts.

**Figure 4.**
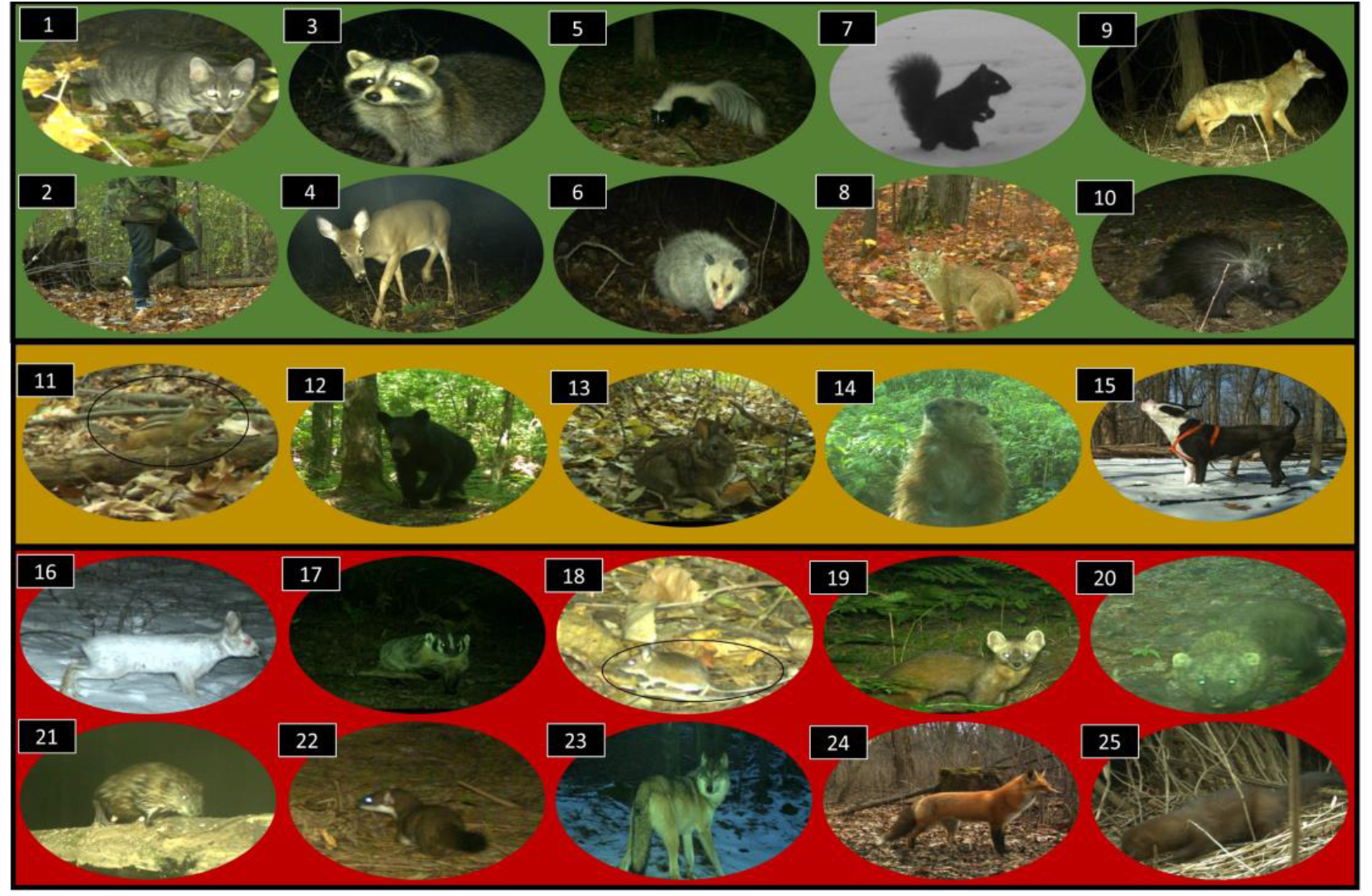
Mammals identified by public volunteers through *Michigan ZoomIN*. Species in green panel obtained ≥90% accuracy: 1) Domestic cat-*Felis catus*, 2) Human*-Homo sapien*, 3) Raccoon*-Procyon lotor*, 4) White-tailed deer*-Odocoileus virginianus*, 5) Striped skunk*-Mephitis mephitis*, 6) Opossum*-Didelphis virginiana*, 7) Squirrel-*Sciurus carolinensis*, 8) Bobcat*-Lynx rufus, 9)* Coyote*-Canis latrans*, 10) Porcupine-*Erethizon dorsatum*; species in yellow panel obtained 80-89% accuracy: 11) Chipmunk-*Tamias striatus*, 12) Black bear-*Ursus americanus*, 13) Cottontail rabbit-*Sylvilagus floridanus*, 14) Groundhog-*Marmota monax*, 15) Domestic dog-*Canis lupus familiaris*; and species in the red panel accuracy < 79% accuracy: 16) Snowshoe hare-*Lepus americanus*, 17) American badger-*Taxidea taxus*, 18) Small mammal-e.g., *Peromyscus maniculatus*, 19) Marten-*Martes americana*, 20) Fisher-*Pekania pennanti*, 21) Muskrat-*Ondatra zibethicus*, 22) Weasel-e.g., *Mustela frenata*, 23) Grey wolf-*Canis lupus*, 24) Red fox-*Vulpes vulpes*, 25) Mink-*Neovison vison*. Gray fox-*Urocyon cinereoargenteus* not shown.

To underscore the importance of considering multiple consensus thresholds, we evaluated results for coyotes as a case study, given their dynamic role within the carnivore community and because they are commonly mistaken for other canid species (Prugh et al. 2009, Colborn et al. 2020). There were 520 classifications of coyotes by users, of which 265 were classified through one of our consensus criteria. Only 9% passed consensus from complete user agreement, while still obtaining >90% accuracy. Instead, the kappa statistic was the best consensus method for identifying this species, comprising 48% of coyote classifications (Figure 3).

We found marginal support for differences in accuracy across species related to study area and species interactions. All study areas had either 99–100% accuracy for common species such as deer, raccoon, and striped skunk. However, rare species, as either being infrequently detected or geographically constrained, had lower accuracy, consistent with our hypothesis. For example, fisher and the small-bodied muskrat were only detected at HMC and SNWR, respectively and had accuracies <50%. Gray wolves (*Canis lupus*) are similar in appearance to coyotes and found only at HMC where they were most commonly misidentified as coyotes. Likewise, though coyotes occurred in every study area, they had the lowest accuracy at HMC. In contrast, the badger (*Taxidea taxus*), which has a noticeably distinct body shape and striking facial pattern, was only captured at UMBS and yet had 75% accuracy.

## DISCUSSION

Our study builds on previous research by suggesting common species, those with distinguishing features, and generally larger species are most accurately identified by the public whereas smaller bodied species and rarer species are less accurately classified. Similarly, Potter et al. (2019) found the most distinct mammal species such as short-beaked echidna (*Tachyglossus aculeatus*) had the highest accuracy from camera surveys in northern Australia, and groups of closely related species with similar morphology, such as mustelids had the most errors. In an African ecosystem, Swanson et al. (2015) found that different species of gazelles as well as weasel-like small mammals were often misidentified from camera data derived from public users. Such variation in accuracy by species may reflect differences in capturing high quality images due to camera placement. As the optimal height for most mesocarnivores is 2-3 feet off the ground (Jacobs and Ausband 2018), smaller or larger bodied species would only be partly captured, which may create more difficulty for users completing identifications.

We also found accuracy varied by study area; though, what specific attributes in geography accounted for these differences is unknown. In addition to habitat characteristics and environmental conditions, the emergent biotic properties such as range shift of mesocarnivores and proximity to urbanization can influence rarity and behavior, altering both the frequency of detection and consequently diminishing user accuracy. Although it is difficult to disentangle the individual components of differences in accuracy between geographies, additional considerations in sampling design could offset differences in study area. For example, additional consideration for standardizing sampling efforts by introducing measuring devices for animal size reference, altering camera height, focusing more on rare species identification in training, and placing multiple cameras at a single location to capture different angles for a species could help account for classification differences related to study area characteristics (Hofmeester et al. 2019). Overall, given the sensitivity of accuracy based upon species and study area underscores the necessity of developing a flexible workflow to process and aggregate user-derived data from camera surveys.

In evaluating our consensus hierarchy, we found that lower threshold consensus criteria such as the third ranking kappa-evenness metric performed nearly as well as 100% agreement. The high accuracy of images classified by these metrics (kappa, kappa and evenness) suggests that hierarchical consensus criteria could be useful in mitigating potential bias from volunteers with varying levels of expertise. By including lower thresholds of kappa and evenness, both of which were more robust to incorrect classifications when aggregating responses, a reliable final classification can still be reached even with multiple individual errors. For researchers, this means that they can be less selective about their volunteer pool and invest less in training and more in data collection or analysis. Given a reasonable mix of expertise in the volunteer pool, more difficult species can still attain sufficient kappa and evenness thresholds from agreement among a smaller number of correct classifications from experienced volunteers. While less experienced members are still valuable in accurately classifying more common species, which may comprise a large proportion of the total images.

Our work highlights the contributions and limitations of untrained participants to accurately classify images of Michigan mammals. We found that most species were eventually correctly identified by employing multiple consensus methods through a robust hierarchical workflow. As a result, we are now confident in lowering the classification per image threshold to below 15 users. Although researchers may find it helpful in future works to first set consensus criteria based on conservative estimates of user knowledge. Future research is needed to determine the degree to which demographic attributes, motivations, environmental stewardship capacity and media content impact classification accuracy and public participation.

## MANAGEMENT IMPLICATIONS

We found collaborating with the public was fruitful for investigating mammals of Michigan– a typical species assemblage found throughout the contiguous United States. We suggest highlighting distinguishing features of species through interactive field guides and species comparisons tools to improve accuracy of user-derived data. Our findings underscore that even with a relaxed criterion of user agreement most species were well represented in the final species classifications. Using a hierarchical workflow with multiple thresholds for reaching consensus to classify images will allow managers to harness a broad volunteer community to maximize workload reduction without compromising accuracy.

## ACKNOWLEDGMENTS

First, we recognize implementing our field research of camera surveys was conducted on lands originally belonging to the People of the Three Fires. We thank the Shiawassee National Wildlife Refuge, the University of Michigan Biological Station, and the Huron Mountain Club for providing access to implement camera surveys for fieldwork on their property. We would like to thank the Applied Wildlife Ecology (AWE) Lab at the University of Michigan for assistance with validating classification. Our work is not human subjects research requiring IRB review, though we remain forever indebted to the nearly 4,000 anonymous volunteers that contributed their time to identify images on *Michigan ZoomIN (MZI)* and the dozens of colleagues that integrated *MZI* into curricula. Our publication used data generated via the Zooniverse.org platform, development of which was funded by generous support including a Global Impact Award from Google and by a grant from the Alfred P. Sloan Foundation. We appreciate technological support from University of Michigan LSA Information Technology and Technology Services teams especially A. Roelofs and P. Knoop. We thank G. Kuhnlein, Michigan News, and the University of Michigan Museum of Natural History for assistance with dissemination on multi-media platforms to recruit volunteers. We appreciate the funding provided by the Detroit Zoological Society, the Huron Mountain Wildlife Foundation, and the University of Michigan Biological Station.

## AUTHOR CONTRIBUTIONS

N.C.H. conceived and designed the research. N.C.H and J.S. developed virtual interface for species identification by public and expert confirmation. G.I.G. and T.C. constructed outreach materials and led trainings on *Michigan ZoomIN*. G.I.G. and R.M. analyzed the data. G.I.G. and N.C.H. wrote the manuscript. All authors contributed to fieldwork, image validation, public engagement and editing the manuscript for final approval.

## Associate Editor

**Summary for online Table of Contents**: Engaging the public proved extremely helpful in processing images of mammals from a large scale camera survey in Michigan, given their high accuracy in species identification. We found that study area, consensus method, and user support best explained accuracy when validating classification from our public science project.

